# SSCM: A method to analyze and predict the pathogenicity of sequence variants

**DOI:** 10.1101/021527

**Authors:** Sharad Vikram, Matthew D. Rasmussen, Eric A. Evans, Imran S. Haque

**Affiliations:** University of California, San Diego; Counsyl

## Abstract

The advent of cost-effective DNA sequencing has provided clinics with high-resolution information about patient’s genetic variants, which has resulted in the need for efficient interpretation of this genomic data. Traditionally, variant interpretation has been dominated by many manual, time-consuming processes due to the disparate forms of relevant information in clinical databases and literature. Computational techniques promise to automate much of this, and while they currently play only a supporting role, their continued improvement for variant interpretation is necessary to tackle the problem of scaling genetic sequencing to ever larger populations. Here, we present SSCM-Pathogenic, a genome-wide, allele-specific score for predicting variant pathogenicity. The score, generated by a semi-supervised clustering algorithm, shows predictive power on clinically relevant mutations, while also displaying predictive ability in noncoding regions of the genome.

## 1 Introduction

It is estimated that 60-70% of medical decision-making is influenced by diagnostic testing and screening [38]. Such testing provides patients with actionable information that allows them to understand their health risks and better plan their future treatment. Accordingly, more informative and available diagnostic testing promises to not only benefit patients, but also improve the efficiency of the health care system overall.

Many clinical screens and diagnostics that have traditionally been based on biochemical testing are today transitioning to a DNA genetic-testing backend. For example, noninvasive prenatal screening (using sequencing to count circulating fetal DNA fragments in a pregnant woman’s bloodstream) is currently supplementing ultrasound-and serum-protein screens for fetal trisomies [15, 10]. Similarly, sequencing-based tests for inborn errors of metabolism are used to screen or diagnose newborns with potentially lethal inherited diseases. The transition towards sequencing-based workflows has been driven economically by the falling cost of sequencing and technically by the high sensitivity and precision of DNA testing compared to noisy protein or mass spectrometry assays [22].

However, the high resolution of sequencing data poses a challenge of *variant interpretation*: it is likely that in each patient, sequencing will reveal new DNA variants, and the clinician must now determine if these newly-observed DNA variants are likely to be pathogenic. These classifications drive all further risk calculations and medical counseling. Current standard methods of variant interpretation [33] are based on a time-consuming, manual integration of multiple data sources, involving extensive database and literature searches, use of computational methods, and multiple rounds of review, taking on average nearly an hour per variant [11, 30]. Frequently, this process does not yield sufficient information, requiring the curator to classify it as a *variant of uncertain significance* (VUS). Depending on the disease, the presence of a VUS may lead a patient to be prescribed additional screening. Naturally, VUS’s can be a source of anxiety for patients who desire concrete results [26, 28]. Due to this additional burden on patients, reducing VUS classifications is a paramount concern.

In theory, computational methods for variant classification could significantly reduce this interpretation burden due to their inherit scalability and objectivity. In fact, the latest guidelines for variant interpretation in clinical sequencing developed by the American College of Medical Genetics and Genomics [33] acknowledge that *in silico* tools can “aid in the interpretation of sequence variants”. However, they also emphasize that *in silico* results are “only predictions” and should not be used as the sole evidence to make a clinical classification. This recommendation is based on the middling accuracy of current computational tools (listed in the guidelines as 65 - 80 % accuracy for missense variants and 60-80% specificity for splicing variants). Improving the accuracy of computational methods is therefore necessary to expand their usability in clinical sequencing.

### 1.1 Computational methods for variant classification

Computational methods typically provide a score per variant or region of the genome, which can then be used to supplement and prioritize the information needed to further classify variants. Broadly speaking, these methods can be divided into several classes based both on their domains of applicability and their biological basis: those that are defined only on coding sequence, those predicting splice sites only, and those defined genome-wide. Typical bases for computation are evolutionary conservation (the use of conservation over multiple species as evidence for functional importance), structure/function (the use of biochemical modeling or inference to predict effects on protein structure), functional inference (the use of functional assays like chromatin accessibility to predict functional regions of the genome), and ensemble techniques (which combine multiple methods to create a more accurate or broader-domained model). A useful list of methods is presented in reference [33].

Thus far, most attention has been on scoring coding variants, particularly missense single nucleotide polymorphisms (SNPs), due their frequency and obvious importance on gene function. Within coding regions, one can use the amino acid translation, reading frame, and similarity to other homologous sequences to gauge how disruptive a variant might be. Many of these features have been heavily used in methods such as SIFT [29] and PolyPhen2 [3] which assign a deleteriousness score based on whether a variant disrupts a significantly conserved region amongst homologous peptide sequences. Recent extensions have also been made for scoring insertion-deletion (indel) variants (PROVEAN [6] and SIFT Indel [18]). Often methods rely on simple probabilistic models to generate scores. LRT calculates how likely a mutation happens given its region [7] and MAPP [37] compares evolutionary variation via the expectation-maximization algorithm for phylogenetics.

Splicing-specific predictive models typically involve statistical learning over experimental splicing data to model the probability that a given mutation will alter the splicing of a transcript. Predicting splicing is particularly important because aberrant splicing can create a very large effect on a downstream protein with a very small nucleotide change (e.g., abrogation of a canonical splice site causing the translation of an extra intron) and because splicing variants can masquerade as silent or small-effect missense variants if interpreted as acting through protein changes. A number of methods have been developed in this area, including MutPred Splice [27], Human Splicing Finder (HSF) [9], MaxEntScan [40], and NNSplice [32]. A major limitation of these methods is the difficulty of predicting noncanonical splice sites, which are often depleted in available training data.

In addition to specific functional impact (missense or splicing), one can also inspect whether a variant disrupts a site that has been conserved, or under negative selection, over long evolutionary time spans. Conservation scores such as GERP [8], PhastCons [35], and PhyloP [31], which predict evolutionary conservation, have been shown to be nearly as powerful as competing methods in predicting deleterious variants [21]. Moreover, these scores can be defined for every base of the genome, enabling genome-wide interpretation of variants. However, used in isolation, evolutionary conservation scores do not take into account a variant’s protein impact and often score regions of the genome, rather than specific alleles.

Interpretation of non-splicing noncoding (intronic or intergenic) variants is made much more challenging by our lack of understanding of the functional impact of these regions of the genome. Newer genome-wide functional assays, such as those performed by the ENCODE and Epigenome Roadmap projects (e.g. chromatin structure, transcription factor binding, and DNA methylation) can provide information about the relative functionality of different regions of the genome [5, 34]. Functional methods such as ChromHMM [12], SegWay [17], and FitCons [16] use this information to predict whether a variant is likely to have functional impact.

Finally, a variety of ensemble methods apply consensus over multiple underlying methods to achieve higher accuracy and broader applicability. For example, the Condel method [14] combines SIFT, PolyPhen-2 and MutationAssessor to better classify missense variation. A particularly interesting recent method is CADD, which combines a large number of scores with a unique training method to achieve high performance [21]. A major challenge in training computational methods (particularly ensemble methods, which may require more data to train each sub-method) is ascertainment bias: “easy” or “obvious” cases are likely to be enriched in databases relative to the entire population of pathogenic variants [39]. CADD avoids this problem by training a classifier to separate known-benign from simulated variants, since both classes can be obtained with little bias. This results in a strong classifier for pathogenicity, likely because the simulated variants (drawn from a realistic distribution of mutation rates without selection) will be enriched for negatively-functional variants versus an observed population (which would be depleted for negatively-selected variants).

Despite the high performance of its ensemble model, CADD’s methodology has a number of downsides. It uses a hand-tuned, very-high-dimensional support vector machine (SVM) to make predictions whose output (distance from separating hyperplane) is not turned into a final score in a straightforward manner. These scores are also difficult to interpret in a probabilistic sense; there is no calibration between a hyperplane distance or CADD score and the probability that a variant is pathogenic. Finally, adding new features into CADD is not straightforward, as the method multiplied features together in a customized manner to account for feature correlation and raise the dimensionality of the data [21].

### 2.2 A new approach

In this work, we present SSCM, Semi-supervised Clustering of Mutations, a fully probabilistic methodology for producing genome-wide, allele-specific variant pathogenicity scores. The key idea is to avoid the need for hard-to-obtain fully labeled training data by training on only partially labeled data. Similar to the ideas introduced by Kircher et al., we use high frequency variants as a partially labeled benign dataset and employ a simulation procedure. However, we view the simulated variants as a mixture of benign and pathogenic (Figure 1b), thus posing the classification problem as semi-supervised clustering. Using this framework, we derive a new classifier, SSCM, and a new variant score, 

~~~
SSCM-Pathogenic
~~~

, that outperforms all of the most popular methods for pathogenicity classification across a wide variety of large, relevant, and realistic datasets. We find, unlike many other scores, that 

~~~
SSCM-Pathogenic
~~~

’s discriminating power extends into many non-coding functional regions, indicating possible future clinical applications. Interestingly, our score not only detects pathogenic variants, but also distinguishes them from otherwise benign tolerated loss-of-function mutations, an important corner case for high specificity. Lastly, our method is interpretable and extensible allowing for additional future improvements. The source code for SSCM is available as open source (https://github.com/counsyl/sscm).

**Figure 1:**
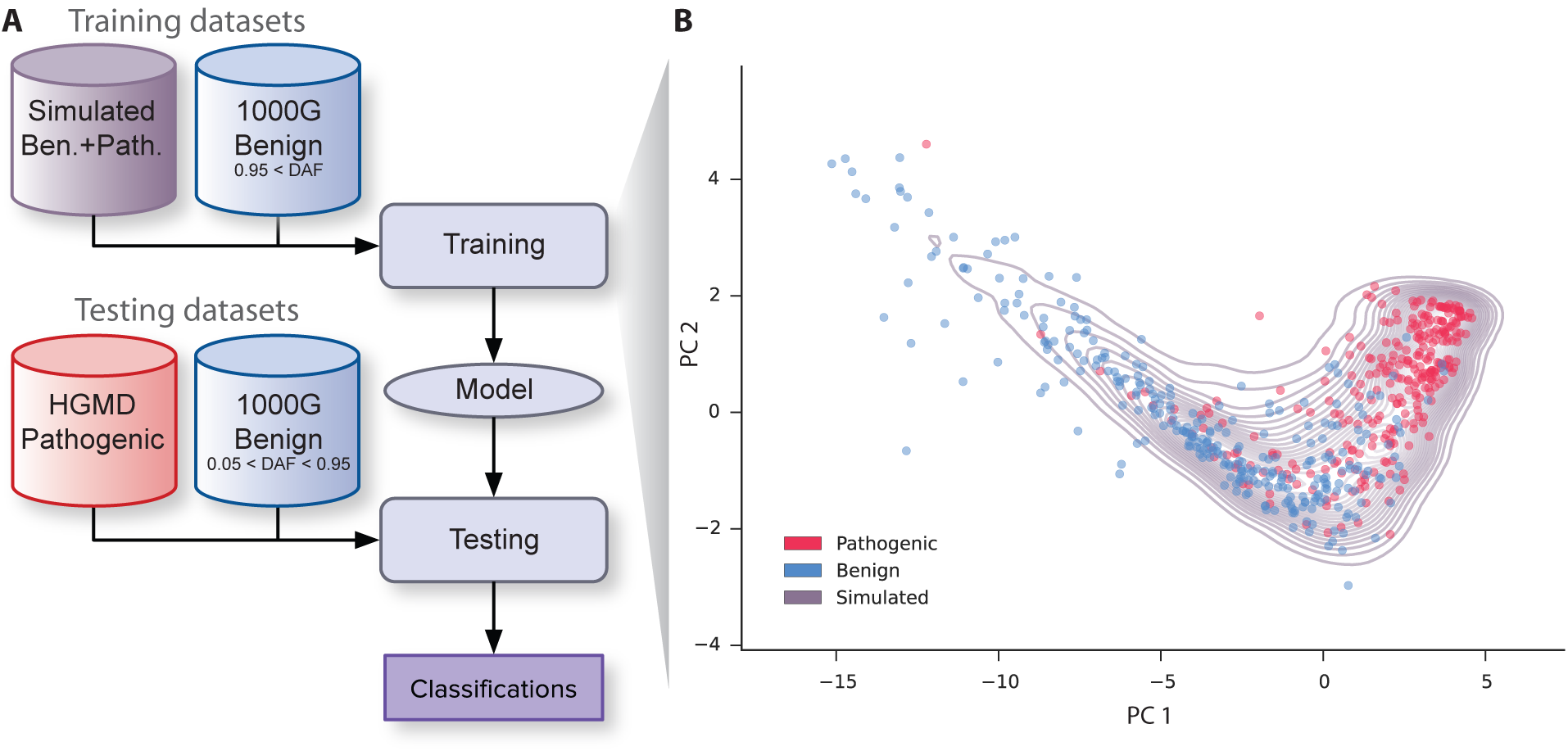
Overview of variant classification training and testing. (A) We trained our model using two datasets: high frequency variants from the 1000 Genomes project, specifically derived allele frequency (DAF) greater than 95%, which are very likely to be benign and randomly simulated variants which are likely to be a mixture of benign and deleterious variants. By treating the simulated variants as unlabeled data, the model learns the distributions of benign and deleterious variants without needing an explicit deleterious training dataset. Classification performance was assessed using distinct test datasets: pathogenic mutations from HGMD and high frequency 1000 Genomes alleles (5% *<* DAF *<* 95%). (B) The top two principle components of the main SSCM features (verPhyloP, verPhastCons, GerpS, SIFT, PolyPhen) were determined for randomly simulated missense variants. A random subset of variants are shown projected into this space from both the benign (blue) and pathogenic (red) test datasets, which are fairly well separated in this feature space. In purple contour lines, a kernel density of the simulated variant distribution is plotted. Notice that it behaves as a mixture of both the deleterious and benign distributions.

## 2 Results

### 2.1 Classification benchmarks

To assess the effectiveness of our approach, we first benchmarked our score, 

~~~
SSCM-Pathogenic
~~~

, against the most successful and popular variant pathogenicity scores, including CADD, SIFT, and PolyPhen2, as well as a purely conservation score, PhyloP. As ground truth, we used pathogenic classifications from the Human Gene Mutation Database (2013.2, Professional Edition) [36] and the ClinVar database Feb 2014 [4]. For benign variants, we filtered 1000 Genomes Project [1] variants by derived allele frequency (*<* 0.95 and *≥* 0.05).

With these benchmarks, we assessed performance across a broad variety of variants. Within coding variants, missense variants are one of the most common and yet difficult to classify. For missense variants, we find that 

~~~
SSCM-Pathogenic
~~~

 shows very strong predictive ability, outperforming the state-of-the-art, CADD, and many other popular protein scores (Figure 2, Figure S1). This increased performance highlights the strength of our learning approach, especially given that these method use many of the same features.

**Figure 2:**
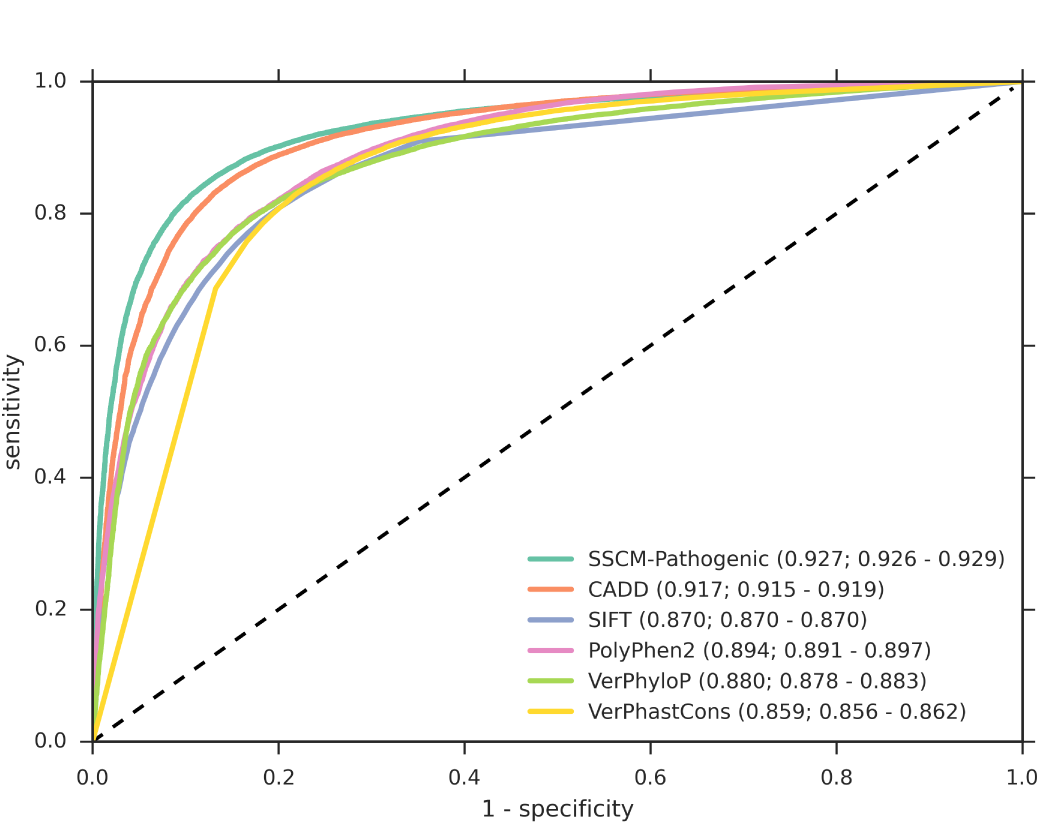
Receiver operator characteristics (ROC) for pathogenic HGMD and benign 1000 Genome missense variants. We obtained pathogenic variants from HGMD (*n* = 63, 363) and benign variants by filtering 1000 Genomes Project variants (*n* = 20, 133) by derived allele frequency (*≥*0.05 and *<* 0.95). SSCM-Pathogenic shows better performance on both datasets. Area-under-the-curve (AUC) values are given along with 95% confidence intervals for the AUCs generated by dataset bootstrap sampling.

For noncanonical splice variants, another difficult yet clinically relevant case, we also find that 

~~~
SSCM-Pathogenic
~~~

 obtains a significantly better receiver operator characteristics (ROC) curve than CADD (Figure 3, Figure S2). This is mostly driven by our inclusion of splicing scores (Figure S4) as features in our model, whereas CADD is relying mostly on conservation scores to classify such variants. Interestingly, we find that 

~~~
SSCM-Pathogenic
~~~

 obtains much higher sensitivity at the lowest false positive rates than all other methods, including the splicing methods. Looking closer, we found that variants classified correctly by 

~~~
SSCM-Pathogenic
~~~

 but missed by splicing methods, tended to be predicted based on their conservation scores, indicating the importance of considering multiple lines of evidence. Notably, 

~~~
SSCM-Pathogenic
~~~

 did not achieve a strictly higher ROC curve, suggesting more could be done to better integrate these features.

**Figure 3:**
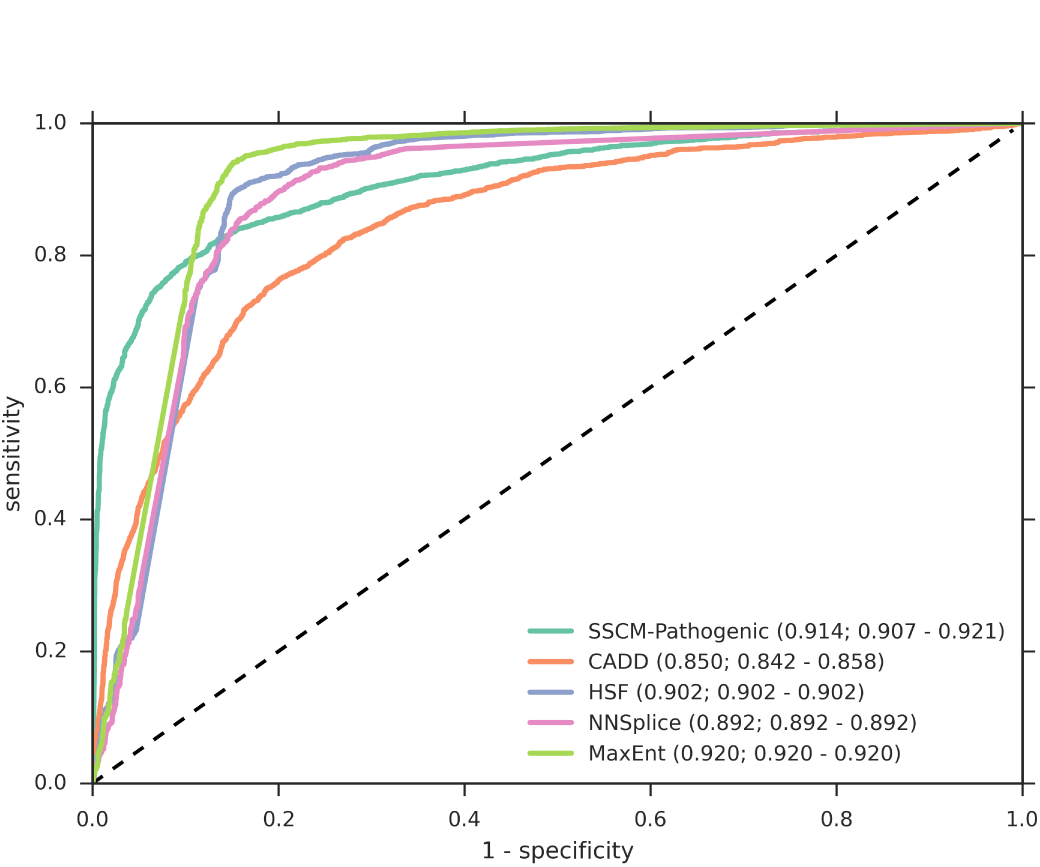
Receiver operator characteristics for pathogenic HGMD and benign 1000 Genome noncanonical splice variants. We obtained pathogenic variants from HGMD (*n* = 2658) and benign variants by filtering 1000 Genomes Project variants (*n* = 6154) by derived allele frequency (*≥*0.05 and *<* 0.95). SSCM-Pathogenic outscores CADD on both datasets while offering better sensitivities for higher specificities than the splice-site scores. Area-under-the-curve (AUC) values are given along with 95% confidence intervals for the AUCs generated by dataset bootstrap sampling.

The results for these methods and variant classes are summarized in Table 1.

**Table 1:**
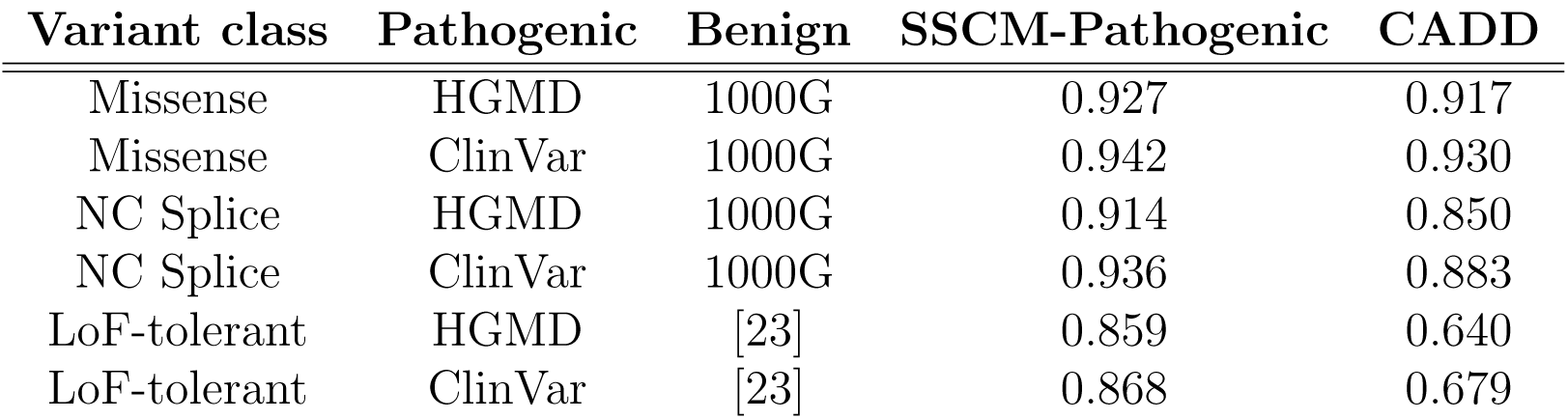
Area-under-the-curve (AUC) values for the receiver operator characteristics of SSCM-Pathogenic and CADD on various variant classes. Shown are results for three different variant classes: missense, noncanonical splice altering (NC splice), and loss of function (LoF) tolerant. Results are fairly consistent across various definitions for benign and pathogenic test datasets. Benign variants (*n* = 7, 633, 050) from the 1000 Genomes Project (1000G) were defined as variants with derived allele frequency *≥*0.05 and *<* 0.95. Benign LoF-tolerated variants (*n* = 228) were obtained from [23]. Pathogenic variants were obtained from HGMD (*n* = 150, 460) and ClinVar (*n* = 47, 007).

### 2.2 Loss-of-Function tolerant mutations

We also benchmarked 

~~~
SSCM-Pathogenic
~~~

 on several loss-of-function (LoF) tolerant variants from MacArthur et al. [23]. Following terminology from MacArthur et al. [24], this class of variants is particularly interesting because although these variants are *damaging*, in that they disrupt a gene’s function, they are not *pathogenic*, so no disease is expressed. Essentially, they are a group of benign variants that can help distinguish between variant scores that merely consider whether a variant is damaging, while ultimately misclassifying the variant’s pathogenicity. We compared 

~~~
SSCM-Pathogenic
~~~

 and CADD in their ability to classify LoF-tolerated variants versus all pathogenic variants from HGMD. We observed significant performance gains for 

~~~
SSCM-Pathogenic
~~~

 (Figure 4), indicating that 

~~~
SSCM-
~~~

Pathogenic is able to better distinguish between pathogenic and damaging variants, a property that is especially important in a clinical setting.

**Figure 4:**
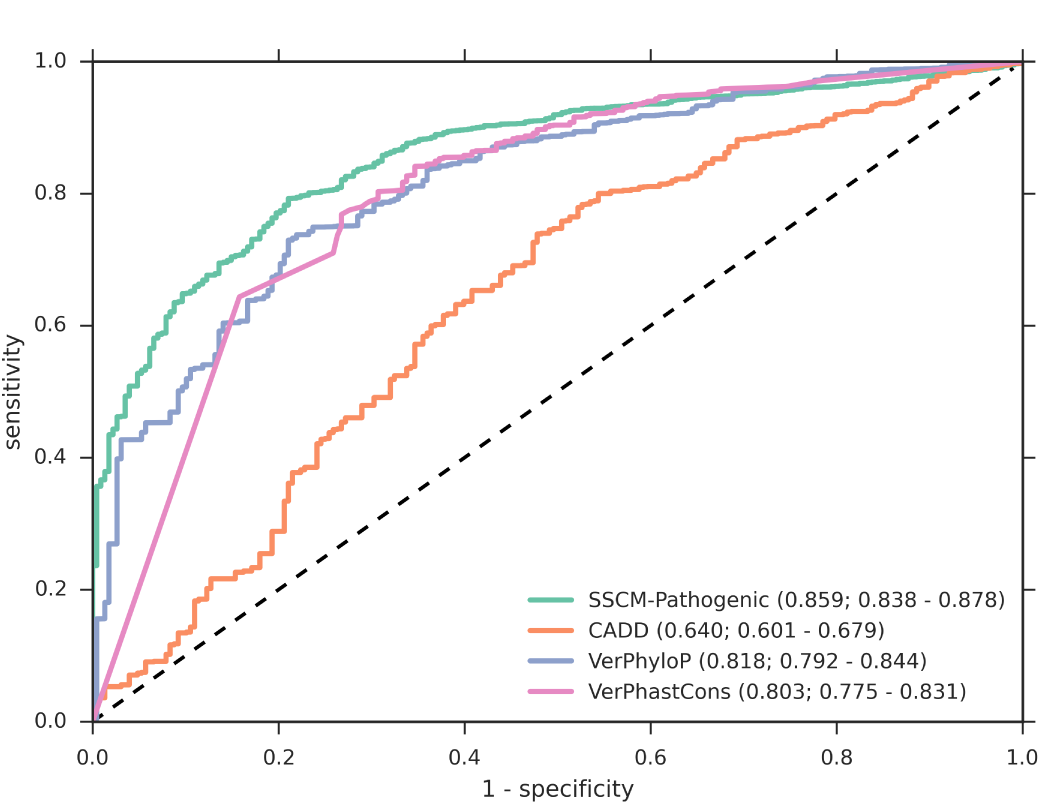
Receiver operator characteristics for pathogenic HGMD and benign LoF-tolerant variants. We obtained pathogenic variants from HGMD (*n* = 150, 460) and MacArthur LoF-tolerant benign variants (*n* = 228). SSCM-Pathogenic shows better performance on both datasets. Area-under-the-curve (AUC) values are given along with 95% confidence intervals for the AUCs generated by dataset bootstrap sampling.

We believe the increased performance comes from 

~~~
SSCM-Pathogenic
~~~

’s ability to better weight conflicting information. For the LoF-tolerant variants, the impact scores (PolyPhen and SIFT) tend favor pathogenicity, while the conservation scores (PhyloP) are in general quite low indicating mutations should be more tolerated. In fact, conservation alone (vertebrate PhyloP) is a fairly good classifier for LoF-tolerant variants (Figure 4).

To further investigate 

~~~
SSCM-Pathogenic
~~~

’s ability to separate pathogenic and damaging mutations, we identified macroscopic genomic trends using our variant-level score. Using LoF-tolerant and recessive genes as defined in MacArthur et al. [23] and autosomal dominant genes from the ClinVar gene database [4], we found that 

~~~
SSCM-Pathogenic
~~~

 is lower on average towards the end of a gene’s transcribed unit, while also finding that different classes showed a spectrum of pathogenicity (Figure 5). The dominant were the most pathogenic on average, followed by recessive and LoF-tolerant. We hypothesized that these results were due conservation features, and showed that vertebrate PhyloP shows the same trend that 

~~~
SSCM-Pathogenic
~~~

 shows in Figure S3. Furthermore, CADD was unable to produce as clear a separation between the gene classes in Figure S3, even though it included the same conservation features.

**Figure 5:**
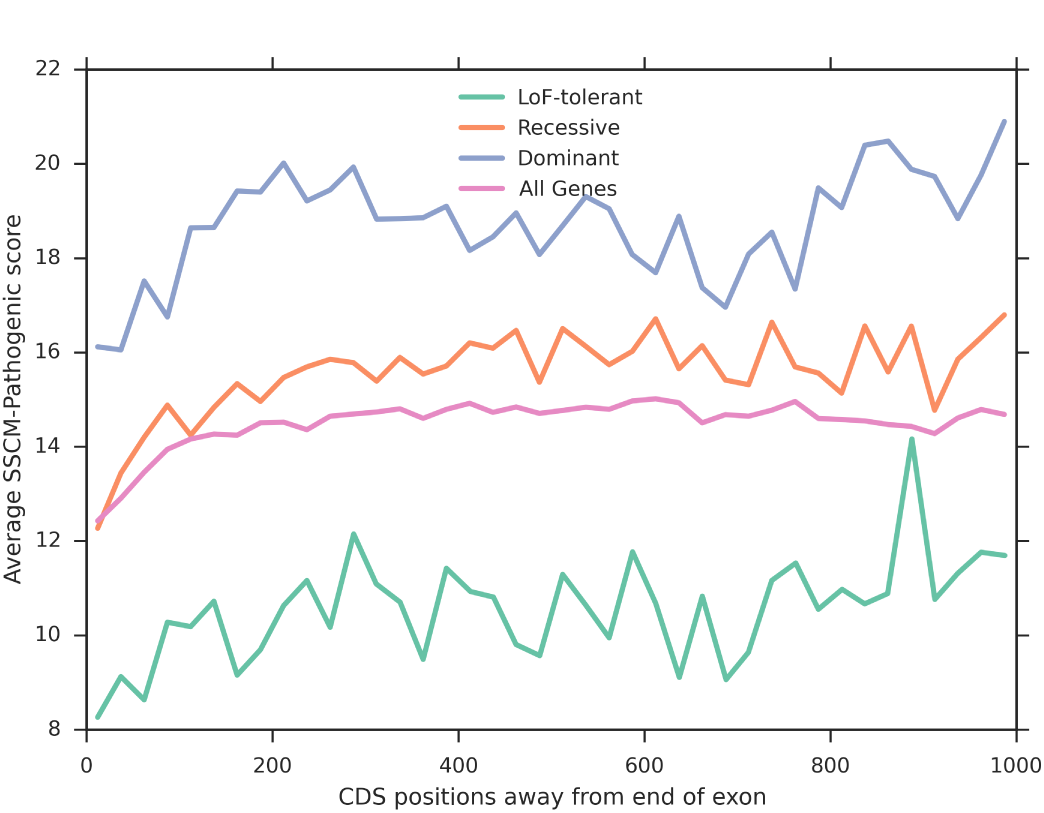
The average SSCM-Pathogenic by coding sequences (CDS) distance away from the end of the exon. We expected variants towards the end of genes to be less pathogenic, and this trend is reflected by SSCM-Pathogenic across a variety of gene classes. Interestingly, the various gene classes show significantly different levels of pathogenicity and they follow the inheritance patterns (LoF-tolerant the least pathogenic and dominate the most).

To better quantify the separation, we generated a gene-level score based on our variant score. For each gene, we computed a new score, LoFA (loss-of-function average), which is the average 

~~~
SSCM-PATHOGENIC
~~~

 score for all stop-gained variants in the gene. Assuming that all stop-gained variants were also loss-of-function, this score would reflect the importance of the gene itself. For example, in a LoF-tolerated gene, all the stop gained mutations should be benign. We found the LoFA for 

~~~
SSCM-PATHOGENIC
~~~

 across the same classes of genes (LoF-tolerant, recessive, and dominant), finding significant separation according to the t-test (Figure 6, Table 2). These results also agree with Khurana et al.’s results with MultiNet [20], which showed the same gene classes can be distinguished with a gene-network based method. CADD, on the other hand, was unable to separate LoF-tolerant from recessive genes nor recessive from dominant genes according to the same set of t-tests. Interestingly, vertebrate PhyloP also passed the same t-tests, again suggesting that conservation metrics are responsible for picking up the difference between damaging and pathogenic variants and that 

~~~
SSCM-PATHOGENIC
~~~

 is capitalizing on conservation better than CADD.

**Table 2:**
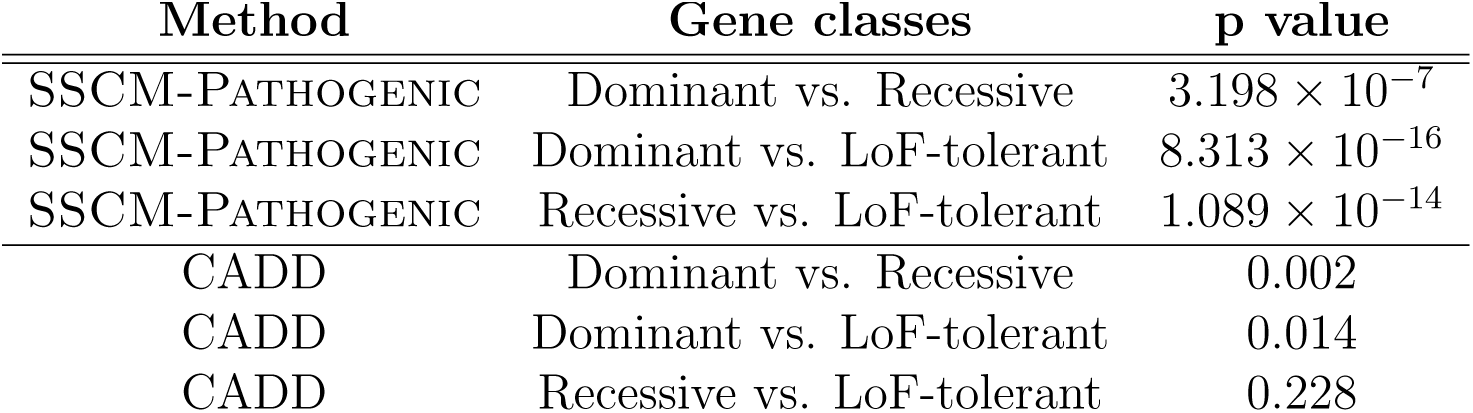
Two-tailed t-test results (*α* = 0.05) for distinguishing between gene classes using “loss-of-function average” (LoFA) for various pathogenicity scores. SSCM-PATHOGENIC is able to successfully separate all three gene classes from each other, indicating SSCM-PATHOGENIC can distinguish between damaging and pathogenic mutations. In contrast CADD is only able to significantly distinguish dominant genes from the other classes.

**Figure 6:**
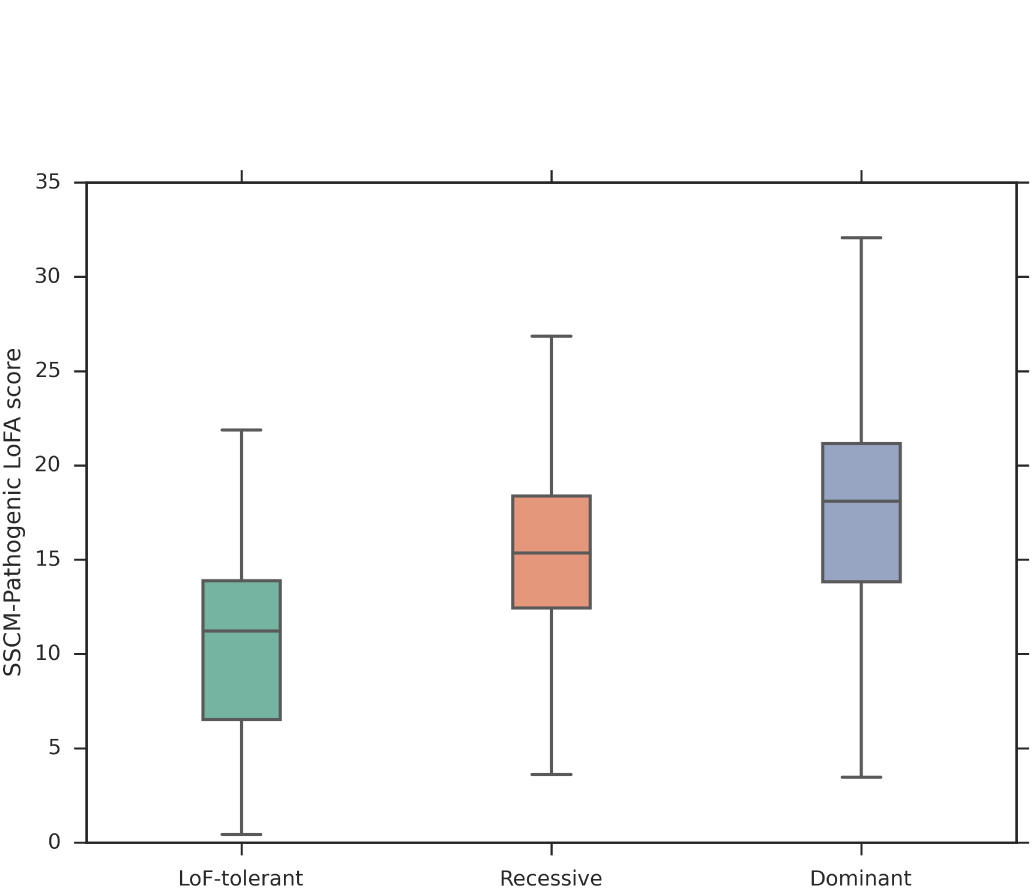
Average SSCM-Pathogenic LoFA scores for three different classes of genes. LoF-tolerant, recessive, and dominant genes were significantly distinguished according to the t-test (*α* = 0.05). The clear separation of LoF-tolerant and recessive genes shows SSCM-PATHOGENIC’s ability to distinguish between damaging and pathogenic mutations, a property shared by conservation metrics like PhyloP, but not CADD.

### 2.3 **Noncoding regions**

Although noncoding region mutations are currently more difficult to interpret relative to missense and splice mutations, we investigated the behavior 

~~~
SSCM-PATHOGENIC
~~~

 for such variants in order better understand the score’s generality. 

~~~
SSCM-PATHOGENIC
~~~

 includes three independent ENcode features (H3K27Ac, H3K4Me3, and H3K4Me1), which we expect to provide the most power in noncoding regions, since these marks are often good predictors of active enhancer and promoter regions. Computing the average 

~~~
SSCM-PATHOGENIC
~~~

 over simulated intronic, intergenic, and untranslated regions (UTRs), we found that 5’ UTRs were enriched for functional elements, resulting in more pathogenicity, followed by 3’ UTRs, intronic, and intergenic regions (Figure 7). These results are largely consistent with those found by Gulko et al. [16], who found that 3’ and 5’ UTRs are more likely to affect the fitness of an organism than intronic and intergenic regions. One difference, however, was that Gulko et al. found that 3’ UTRs are more likely to affect fitness than 5’ UTRs.

**Figure 7:**
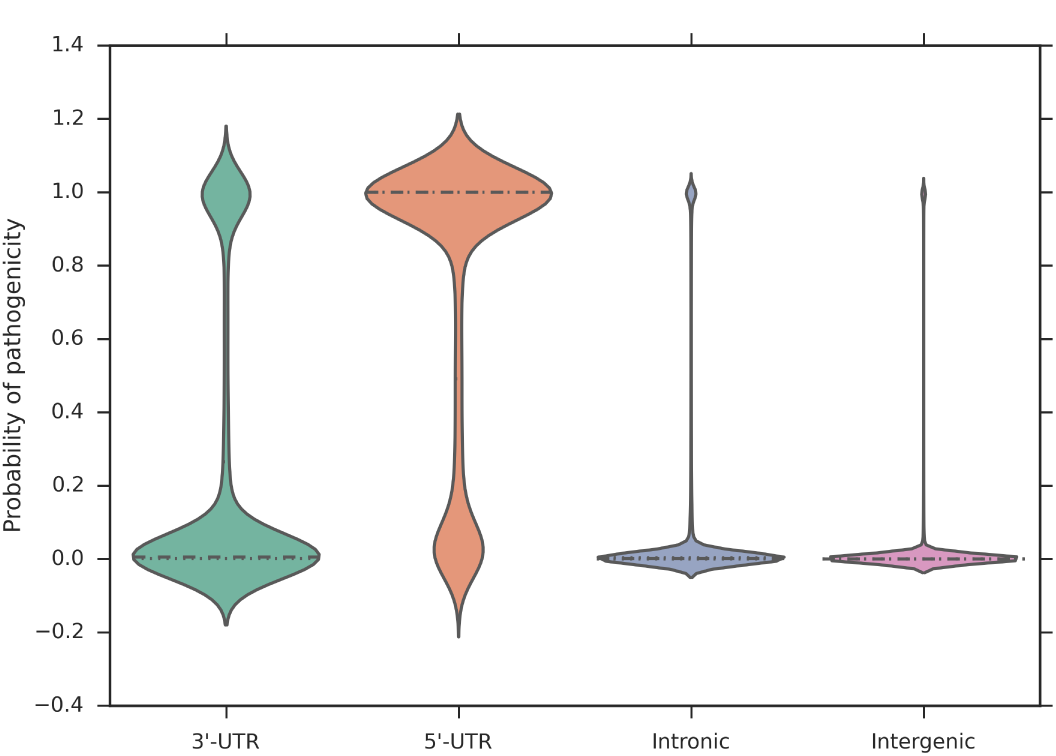
Distribution of SSCM-Pathogenic over different noncoding regions. SSCM-PATHOGENIC shows a clear difference among UTRs, intronic, and intergenic regions and supports results that the 5’ UTRs are enriched whereas intronic and intergenic regions are depleted for pathogenicity. Note that all values are within [0, 1] even though the density curve extends slightly outside these bounds.

### 2.4 Comparison to supervised model

We also compared our methodology against a simple, supervised learning approach. We fitted a model using the same features, except we performed a maximum likelihood fit to HGMD pathogenic mutations and 1000 Genomes benign mutations, rather than clustering partially-labeled simulated data. This model performs marginally better than 

~~~
SSCM-PATHOGENIC
~~~

 on ClinVar missense and splice mutations (Figure S5), which was expected given the overall similarity between mutations in ClinVar and HGMD, resulting in test data very similar to the training. However, on LoF-tolerant mutations, 

~~~
SSCM-PATHOGENIC
~~~

 slightly outperformed the supervised model (Figure S6). Further examining the supervised model revealed the distributions it found had lower variance and its scores tended to be more extreme, which is typical of overfitting.

## 3 Materials and Methods

### 3.1 Training data and features

The first step in our statistical learning method was to obtain data. Our training datasets mimicked those used by Kircher et al. First we defined a “known benign” data set by filtering variants from the 1000 Genomes Project [2] by high derived allele frequency (*≥* 0.95), as we assumed that alleles with extremely high frequency were benign, resulting in a set of 881,924 SNPs. Next, we generated a “simulated” data set of 1,405,358 variants using CADD’s variant simulation software (http://cadd.gs.washington.edu/static/NG-TR35288_Supp_File1_simulator.zip, downloaded Feb 9, 2014). The program mutates a locus according to local mutation rates in a sliding 1.1Mb window. These local mutation rates were obtained by comparing the human genome to an inferred human-chimpanzee ancestor and bases were changed according to a genome wide-determined substitution matrix. For all analyses, we used the same simulation parameters as listed in the supplement of [21]. See Figure 1a for an overview of the training and testing datasets and workflow.

All variants were annotated with features from Ensembl’s Variant Effect Predictor version 68 [25]. These annotations cover a wide range of scores, from conservation features (PhastCons, phyloP, GERP++, etc.) [35, 31, 8] and missense variant scores (SIFT, PolyPhen2) [29, 3] to an array of regulatory scores (ENcode) [5]. We added three splice site features to the datasets, namely HSF, NNSplice, and MaxEnt, provided by Interactive Biosoftware’s Alamut Batch v1.1.11 [9, 32, 40, 19].

Although VEP provides 63 annotations for each variant, our final model only included 12 features total (Table 3). Many features were initially not included because they had no immediate tie to pathogenicity (e.g. GC count). To select out of the remaining features, we first allocated a portion of the HGMD and ClinVar pathogenic variants and a portion of the 1000 Genomes benign variants into a validation set. We chose the set of features that maximized the validation score, resulting in a set of 9 features from the original 63 annotations, plus the three additional splice features.

**Table 3:**
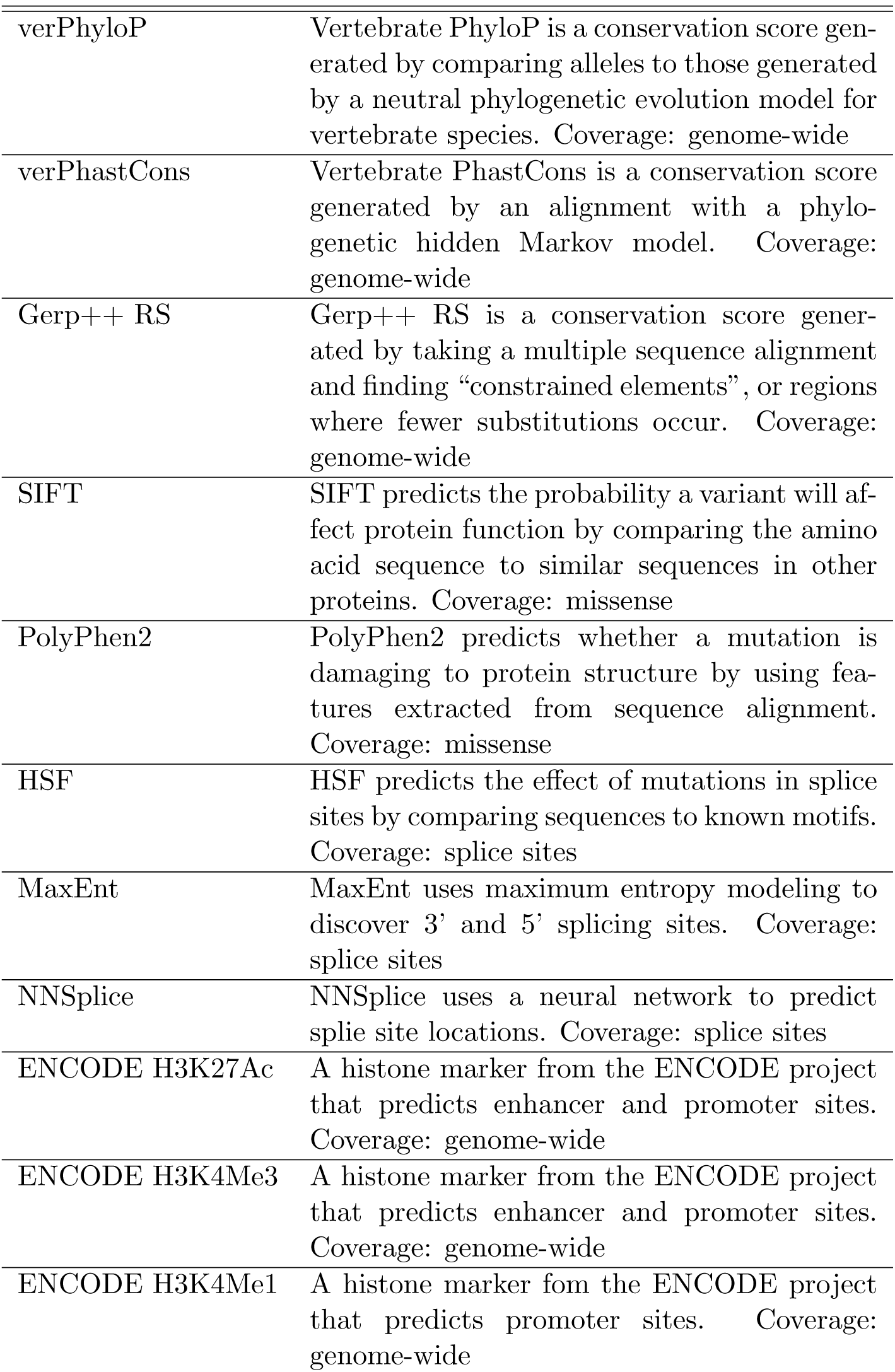
List of all the features used in our method that resulted in the largest validation accuracy.

### 3.2 Generative model for mutations

We first designed a generative model for the simulated dataset, specified as follows. Let 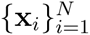 represent the simulated variants. We assume two clusters in the data: pathogenic and benign, and then assume a hidden variable *z*_*i*_ which represents a variant’s assignment to either the pathogenic cluster or benign cluster. Each variant has a set of *D* features associated with it, 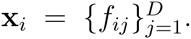 Features for a variant, which could be either vector or scalar, are conditionally independent given the cluster assignment *z*_*i*_ and each have a specific distribution drawing its parameters from the parameter matrix *θ*, *p*_*j*_(*f*_*ij*_|θ_*z*_*i,j*). We also assumed a multinomial distribution on *z*_*i*_ with parameter ***π*** with a Dirichlet prior on ***π*** with hyperparameter *α*. This generative model is pictured in Figure 8:

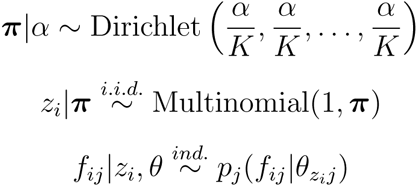

**Figure 8:**
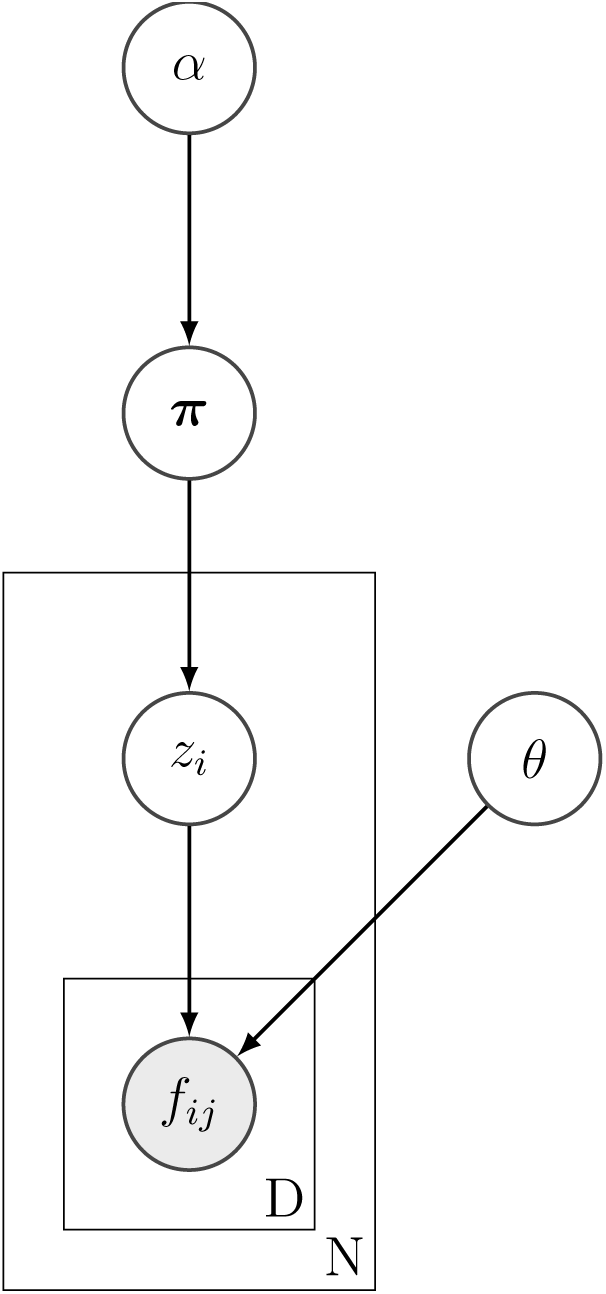
A generative model for a simulated variant dataset with independent features. Conditioned on its assigned cluster *z*_*i*_ (e.g. benign, deleterious), a variant *i* has several independent features *f*_*ij*_ (e.g. conservation, amino-acid features, functional scores). These features each have their own distribution, either a multinomial or multivariate Gaussian, which combined have parameters *θ*. Cluster assignments are modeled with a multinomial prior of mixing weights *π*, which in turn has a Dirichlet prior with hyperparameter *α*.

We assigned univariate Gaussian or multinomial distributions to each of the *D* features. We found these distributions to be both convenient To mitigate the effect of the naive Bayes assumption, we allowed grouping features into vectors and assigning a multivariate Gaussian distribution to the compound feature vector. To find clusters in this generative model, we used the expectation-maximization algorithm to estimate the parameters ***π*** and *θ*. EM iteratively calculates posterior probabilities of the hidden variable *z*_*i*_ for each variant and then updates the values of the parameters ***π*** and *θ* to maximize the likelihood of the data given the soft assignments of *z*_*i*_.

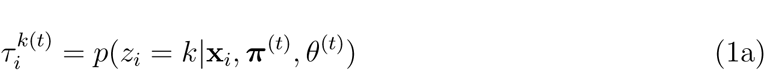

updates for the parameter ***π*** = [*π*_1_, *π*_2_, *…, π_K_*] were:

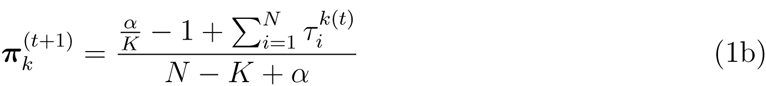

For a Gaussian feature for the cluster assignment *z*_*i*_ = *a* and feature *j* = *b*, the updates are:

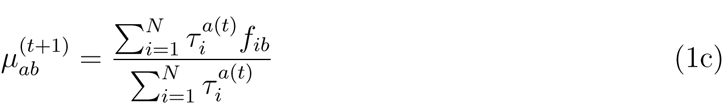

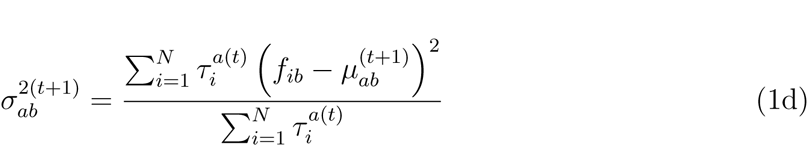

For a multinomial feature for the cluster assignment *z*_*i*_ = *a* and feature *j* = *b*, the updates for each component *v* of the parameter vector *p*_*ab*_ = [*p*_*ab*0_, *p*_*ab1*_,…, *p*_*abL*_] are:

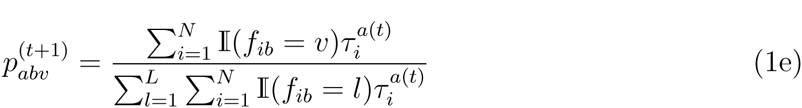

For a multivariate Gaussian feature for the cluster assignment *z*_*i*_ = *a* and feature *j* = *b*, the updates are:

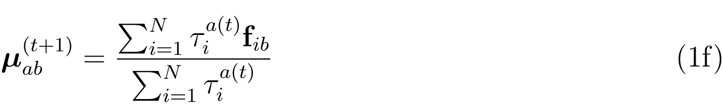

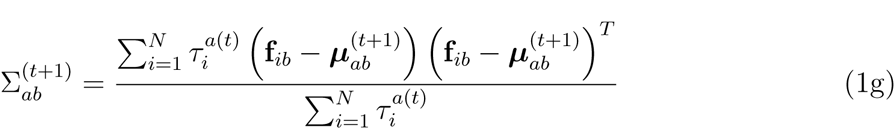

To incorporate the labeled known benign dataset into the algorithm, we obtained maximum likelihood estimates of the parameters from the data and initialized the parameters of the first cluster in the EM to these estimates. We also held the parameters of the benign cluster constant throughout the algorithm, allowing only the pathogenic cluster’s parameters to be updated on every iteration.

We ran EM until convergence, a process that took about 8.5 hours on a single core of a 2.9GHz Linux server. Multiple random initializations of the parameters ended up with the same set of parameter values, implying that EM terminated at a reasonable local maximum. Scores for unknown variants were generated by calculating the posterior probability of assignment to each of the clusters; for interpretability and comparability, we often used the negative posterior log probability of belonging to the benign cluster to match the scale of CADD.

### 3.3 Handling Missing Data

Both the simulated dataset and the known benign dataset had large amounts of missing data; over 50% of values in the files were N/A. This was largely due to features being defined only in certain regions of the genome. For example, SIFT is only defined on missense variants whereas PhyloP and PhastCons are defined on a majority of the genome. To account for these missing values in a Bayesian manner, we integrated out the features that were not present in a particular variant. Due to the naive Bayes assumption, calculating the posterior probability of a mutation belonging to a cluster only involved using likelihoods of features present at a locus. We believe this was a reasonable way to treat the missing data, as the probabilistic model calculated a posterior probability conditioned on only the data available.

We also modified the updates for the multivariate Gaussian parameters by calculating the mean vector and covariance matrix on a feature by feature basis, rather than in a vectorized manner to handle missing data. Due to the missing data, there was the possibility of a non-positive semidefinite covariance matrix, which we corrected by computing the eigendecomposition, setting the negative eigenvalues to a slightly positive number, and regenerating the matrix. However, this problem only occurred when there were very small amounts of data.

## 4 Discussion

The growing use of genome sequencing in the clinic has presented many challenges, one of the most prominent being the need to accurately and thoroughly interpret a patient’s genetic variants and their influence on disease status or risk. While many other aspects of genomic testing are undergoing dramatic improvements in efficiency and performance, current approaches to variant classification are manually intensive and time-consuming, thus creating an “interpretation bottleneck” within the overall genomic work flow. Motivated by this challenge, we have introduced a new computational method for classifying variant pathogenicity, which we have demonstrated out-performs all of the most popular current approaches.

We have obtained these improvements by carefully posing the problem as semi-supervised learning, which avoided the long-standing challenge of obtaining unbiased training data. While there are large, growing databases of variant classifications, such as HGMD and ClinVar, these datasets are still dominated by fairly obvious cases. Accordingly, we have found that directly training on such data overfits on variants for which classification is already easiest (Figure S6). While it has been possible for quite sometime to obtain comprehensive benign variant examples by conditioning on high allele frequency in public databases, such as 1000 Genomes [1] and the Exome Sequencing Project (ESP) [13], comprehensive pathogenic variants have been hard to come by. We overcame this challenge by simulating variants across the genome using a model of realistic mutation rates. This produced a distribution of variants in the absence of natural selection, thus enriching for pathogenic variants. We found that this simulated distribution in combination with a labeled benign dataset provided enough information to learn a classifier for pathogenicity that showed far less bias than a fully supervised model.

To better understand our performance gains, we inspected the power of each of our features. Overall, we found that evolutionary conservation consistently contributed to our score’s performance. This was the case in distinguishing merely damaging loss-of-function variants from pathogenic (Figure 4) and in general trends such as the depletion of pathogenic truncating variants from the 3’-end of genes (Figure S3). This is consistent with the observation that conservation plays a significant role in the CADD method’s performance [16]. However, in the case of 

~~~
SSCM-PATHOGENIC
~~~

 there are many instances where other features play a more important role. For missense mutations, SSCM-Pathogenic benefits from missense-specific features such as SIFT and PolyPhen2 and outperforms the pure conservation score PhyloP (Figure 2) and in intronic and intergenic regions benefits from ENcode features (Figure 7).

Although, we have demonstrated clear performance gains, this work still only provides one piece of the total evidence needed to properly classify a variant in a clinical setting [33]. Going forward, additional work will be needed to fully realize the potential for computational methods to address the interpretation bottleneck that exists in current genomic testing.

A major benefit of our parametric generative model is its simplicity and interpretability. However, given that this approach uses simulated data, large amounts of data can be obtained, and thus non-parametric techniques are likely feasible. For example, approaches such as Dirichlet-process mixture models or kernel density estimation may be able to better capture the complex boundaries between benign and pathogenic clusters.

Although we classified two clusters in this work (benign and pathogenic) there are signs that multiple distinct clusters may be present (Figure 1, Figure S7, Figure S8). For example, there may be multiple kinds of pathogenic variants, each with their own characteristics. With unsupervised learning techniques, we could discover new classes of variants or learn more about known variants.

Two important aspects of SSCM are its reproducibility and extensibility. The choice of datasets to train and test on is of utmost importance and all projects should be open about these decisions to avoid overlapping datasets in the process. To help with this aspect, we are open-sourcing our method and have been explicit about all training and testing datasets. We also aimed to make our score as extensible as possible. Our naive Bayes assumption and treatment of missing data allows for any annotation to be a part of the process. This enables future scores to be added with ease.

## 5 Supplementary Material

**Figure S1:**
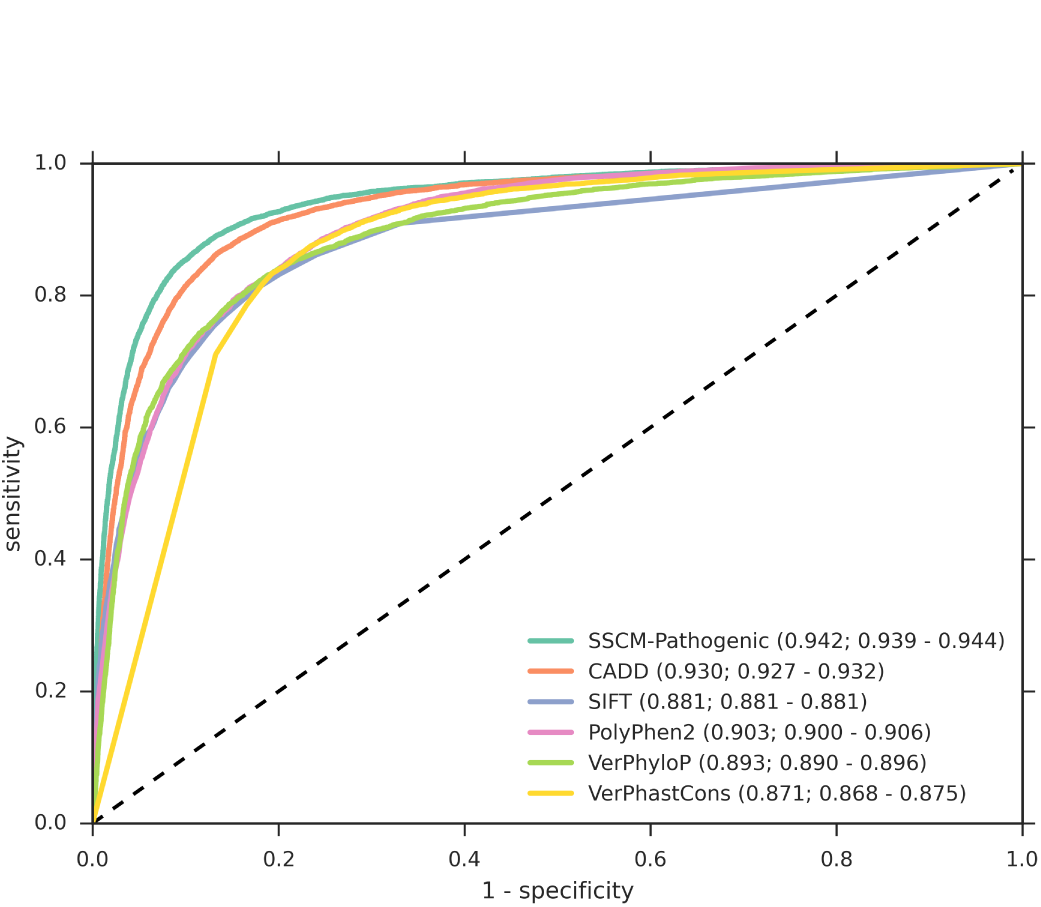
Receiver operator characteristics (ROC) for pathogenic ClinVar and benign 1000 Genomes missense variants. We obtained pathogenic variants from ClinVar (*n* = 18783) and benign variants by filtering 1000 Genomes Project variants (*n* = 20133) by derived allele frequency (0.05 *≤* DAF *<* 0.95). Area-under-the-curve (AUC) values are given along with 95% confidence intervals for the AUCs generated by dataset bootstrap sampling.

**Figure S2:**
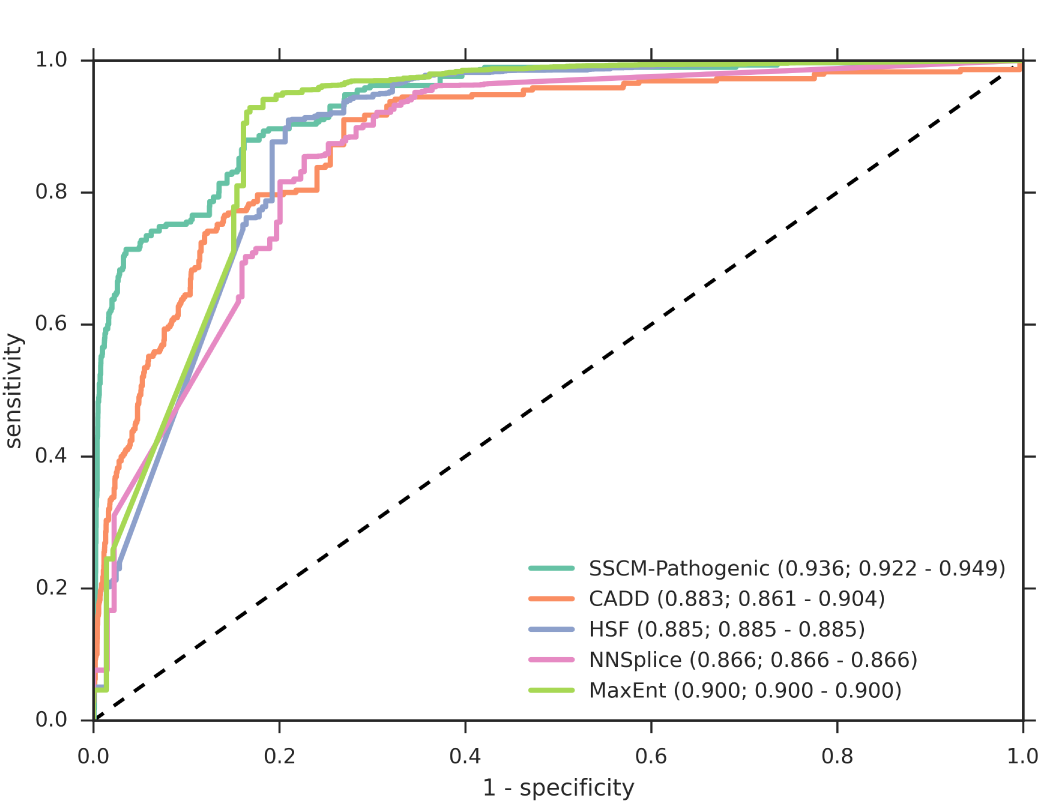
Receiver operator characteristics (ROC) for pathogenic ClinVar and benign 1000 Genomes noncanonical splice variants. We obtained pathogenic variants from ClinVar (*n* = 290) and benign variants by filtering 1000 Genomes Project variants (*n* = 6158) by derived allele frequency (0.05 *≤* DAF *<* 0.95). Area-under-the-curve (AUC) values are given along with 95% confidence intervals for the AUCs generated by dataset bootstrap sampling.

**Figure S3:**
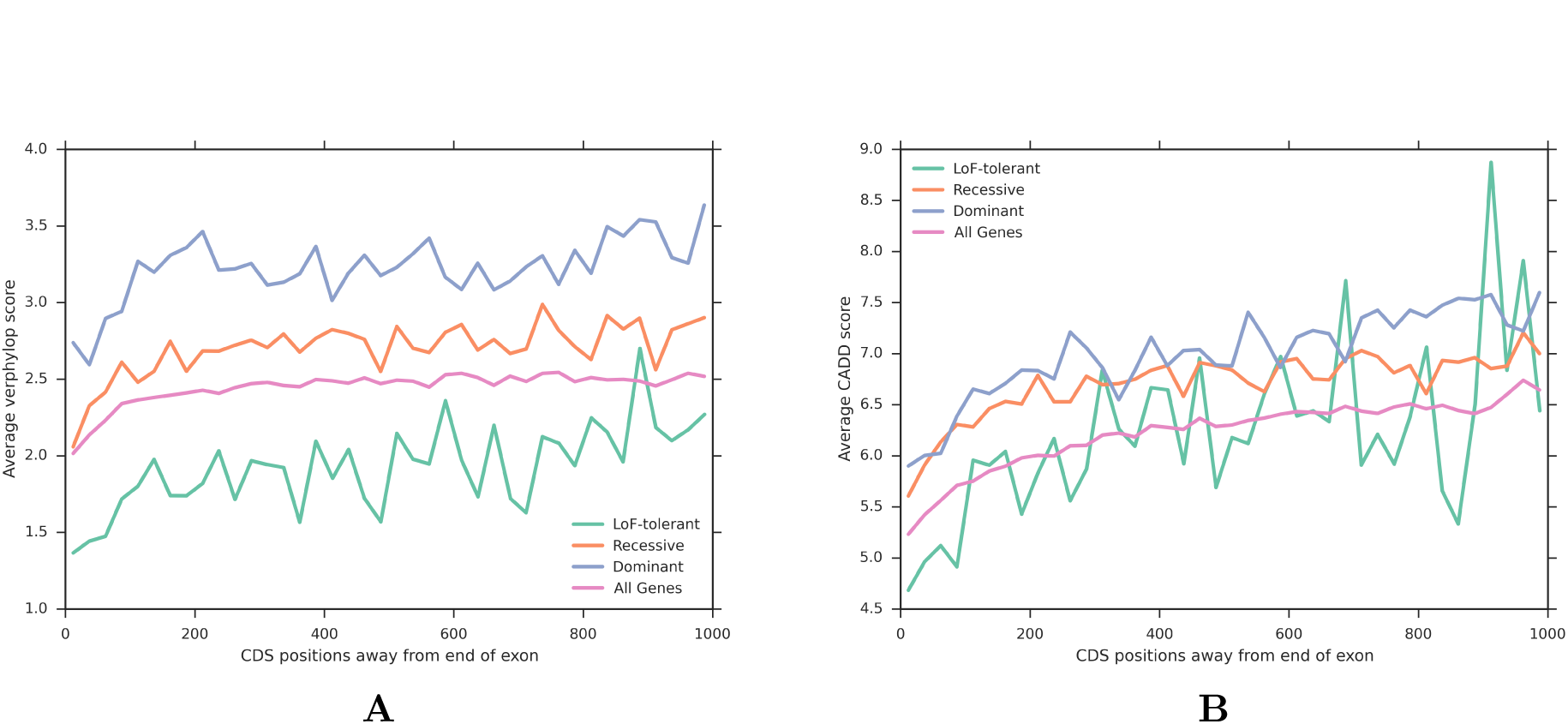
The average vertebrate PhyloP and CADD score in terms of coding (CDS) distance upstream of the gene’s stop codon. (A) Conservation appears to explain much of SSCM-PATHOGENIC’s ability to identify macroscopic gene trends (see Figure 5). (B) In contrast, CADD does not as clearly distinguish such genes either overall or along the length of the coding sequence.

**Figure S4:**
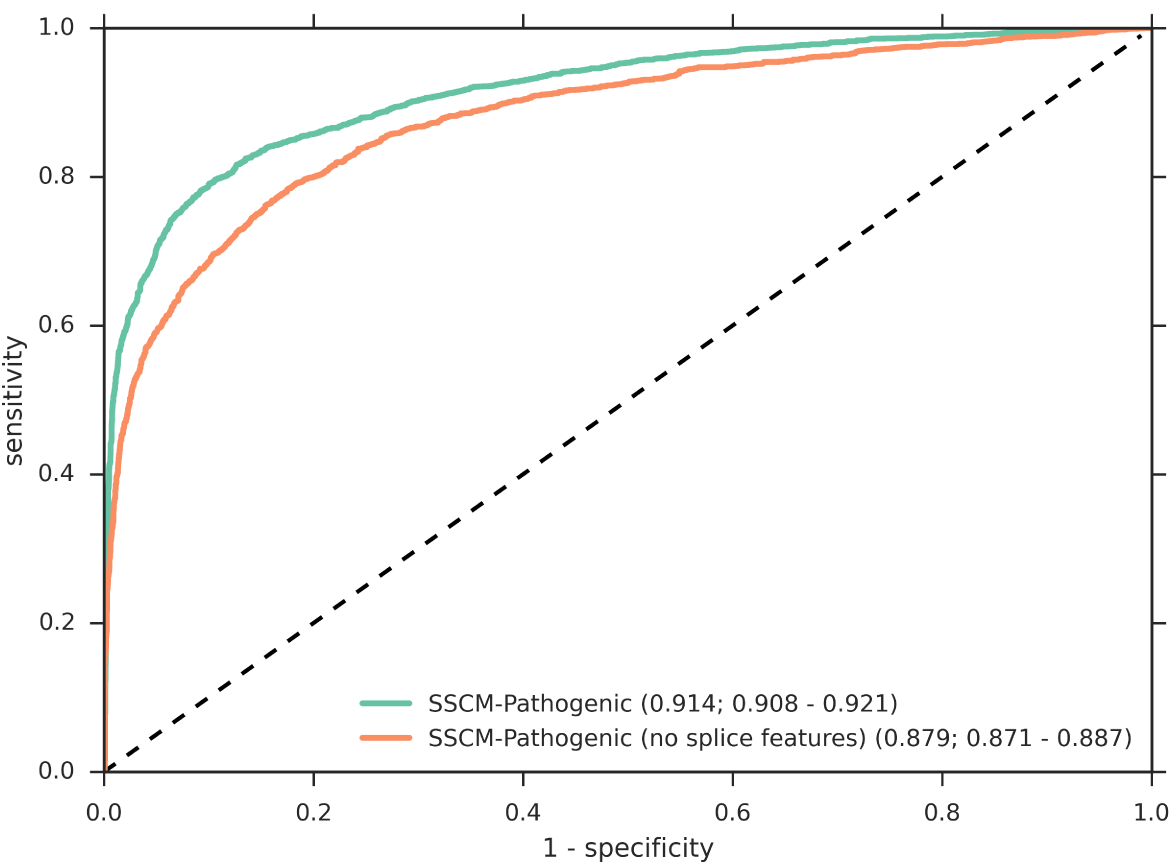
Receiver operator characteristics (ROC) for pathogenic HGMD and benign 1000 Genomes noncanonical splice variants. We obtained pathogenic variants from HGMD (*n* = 2658) and benign variants by filtering 1000 Genomes Project variants (*n* = 6154) by derived allele frequency (0.05 *≤* DAF *<* 0.95). This particular ROC shows the same model used with and without the inclusion of splice features (HSF, MaxEntScan, NNSplice). In this scenario, the inclusion of splice features increases performance. Area-under-the-curve (AUC) values are given along with 95% confidence intervals for the AUCs generated by dataset bootstrap sampling.

**Figure S5:**
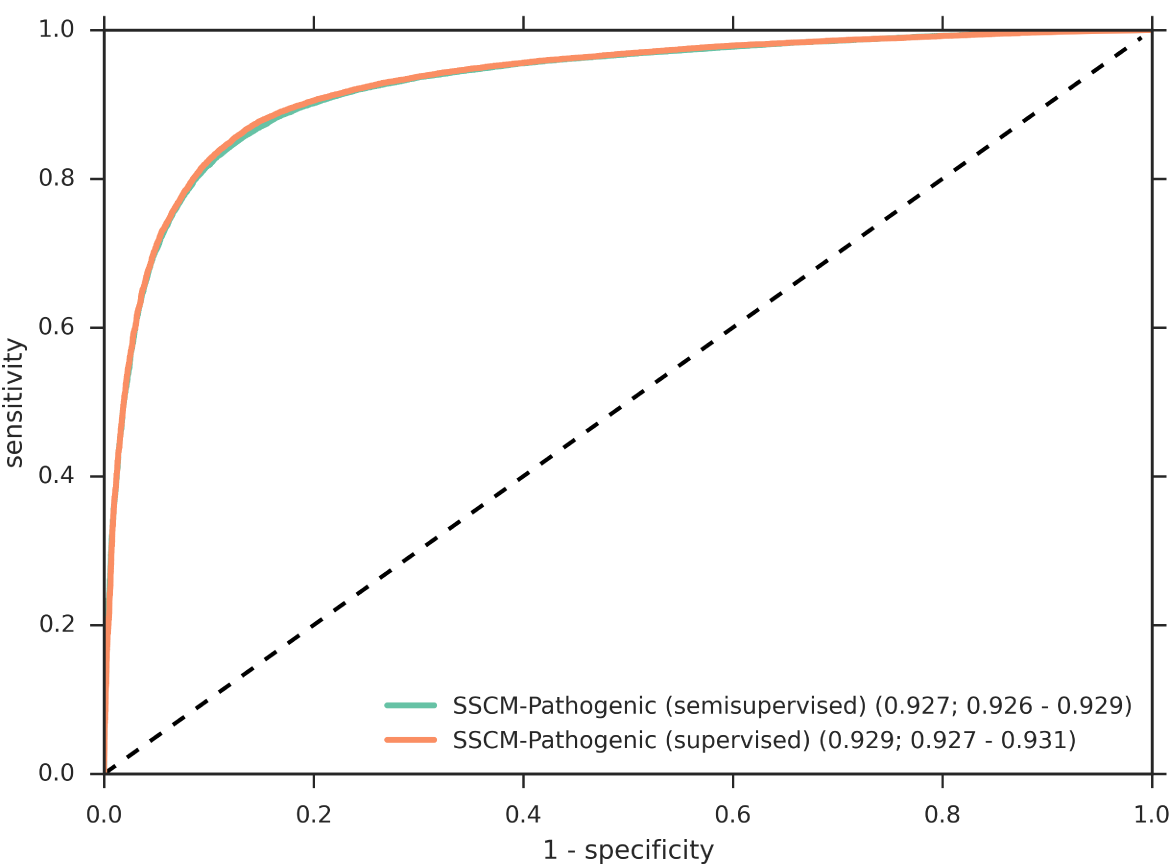
Receiver operator characteristics (ROC) for pathogenic HGMD and benign 1000 Genomes missense variants. We obtained pathogenic variants from HGMD Pro (*n* = 63363) and benign variants by filtering 1000 Genomes Project variants (*n* = 20133) by derived allele frequency(*≥* 0.05 and *<* 0.95). SSCM-PATHOGENIC shows better performance on both datasets. This ROC compares models trained with semisupervised learning and with supervised learning. Area-under-the-curve (AUC) values are printed along with 95% confidence intervals for the AUCs generated by dataset bootstrap sampling.

**Figure S6:**
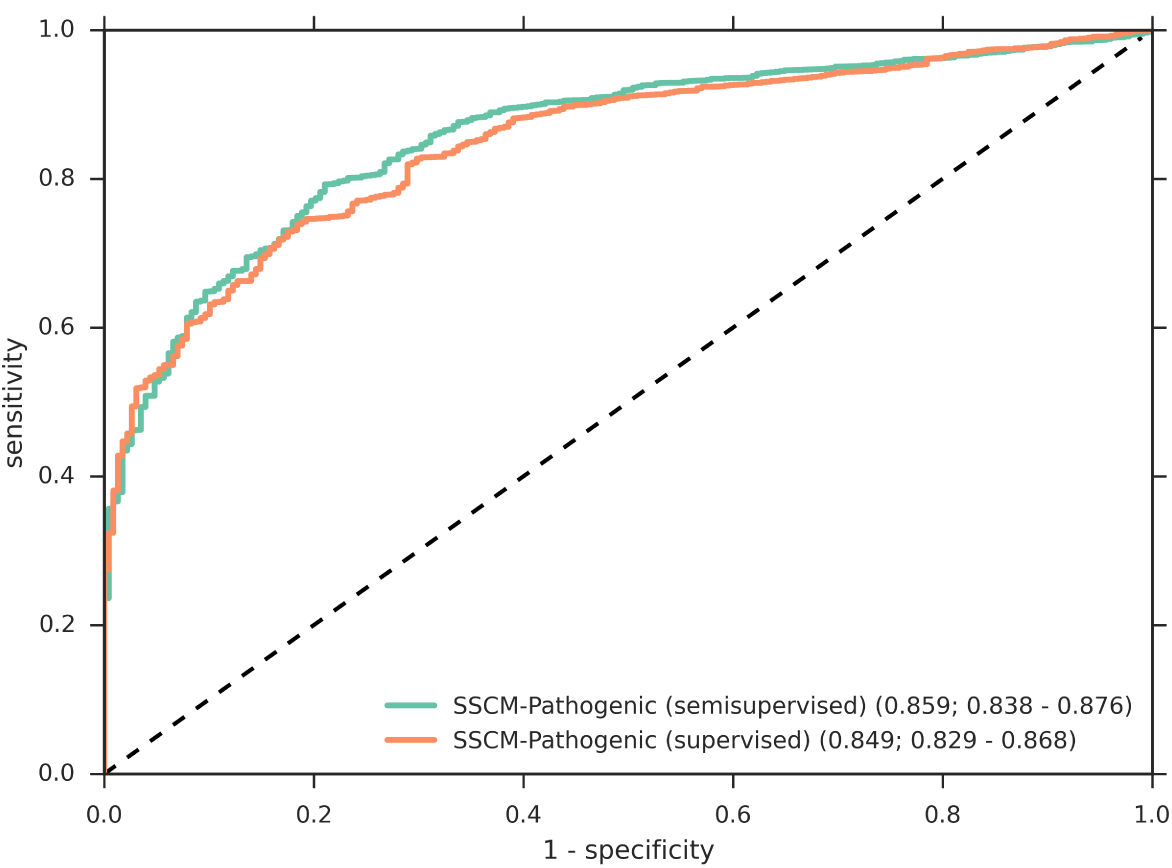
Receiver operator characteristics (ROC) for a LoF-tolerant mutations with semi-supervised learning model and a supervised learning model. We obtained pathogenic variants from HGMD (*n* = 150460) and MacArthur LoF-tolerant benign variants (*n* = 228). This particular ROC shows the same model trained with semi-supervised learning and with supervised learning. In this scenario, the semi-supervised model performs marginally better. Area-under-the-curve (AUC) values are given along with 95% confidence intervals for the AUCs generated by dataset bootstrap sampling.

**Figure S7:**
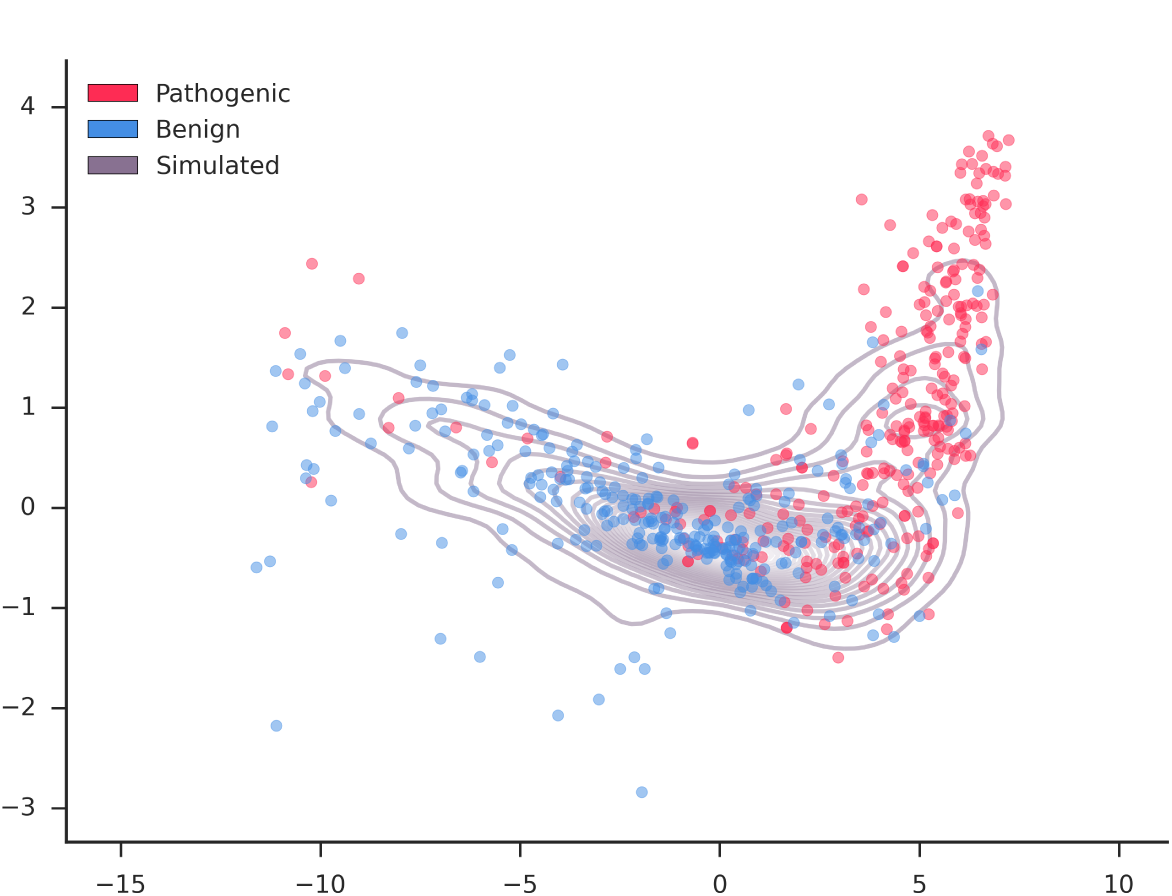
PCA plot of NC splice mutations. The top two principal components of the main SSCM features (verPhyloP, verPhastCons, hsf, GerpS, MaxEntScan, NNSplice) were determined for randomly simulated NC splice variants. A random subset of variants are shown projected into this space from both the benign (blue) and pathogenic (red) test datasets. In purple contour lines, a kernel density of the simulated variant distribution is plotted.

**Figure S8:**
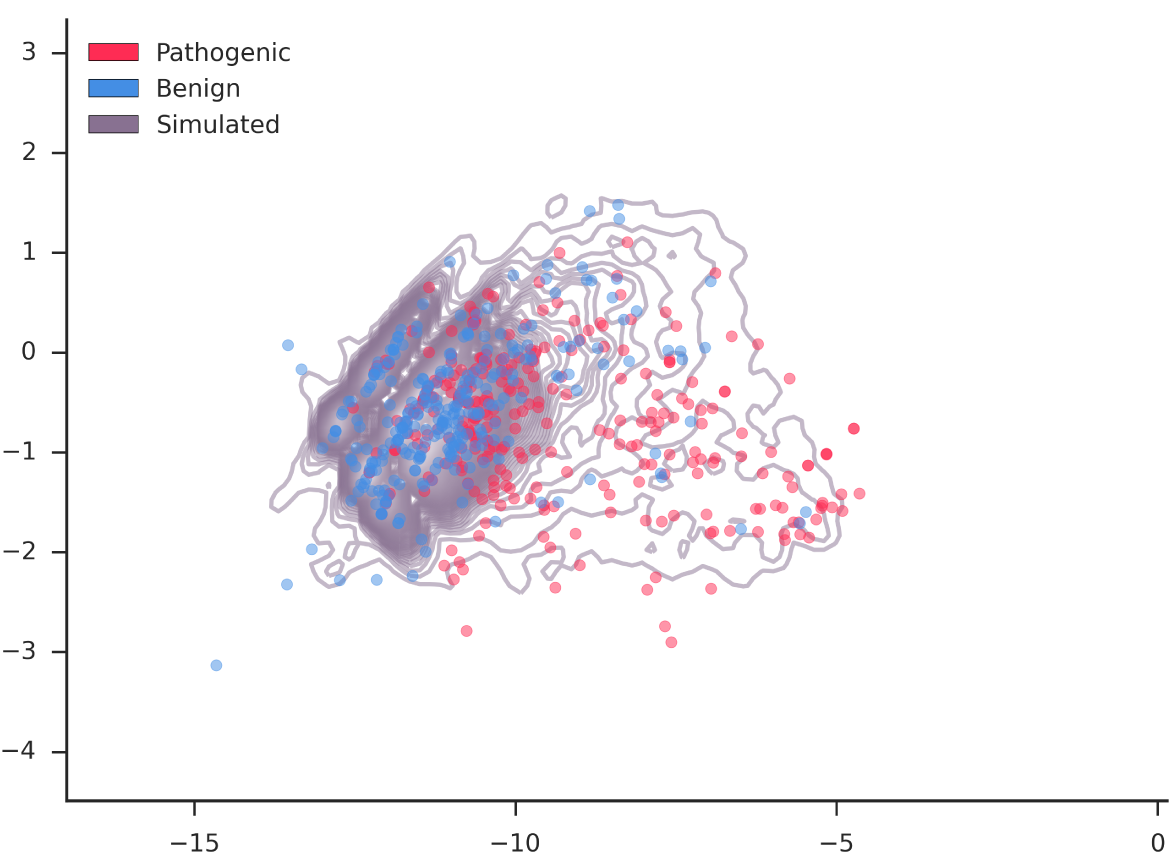
PCA plot of noncoding-region mutations. The top two principle components of the main SSCM features (verPhyloP, verPhastCons, GerpS, ENcode H3K27Ac, ENcode H3K4Me3, ENcode H3K4Me1) were determined for randomly simulated intergenic, regulatory and intronic variants. A random subset of variants are shown projected into this space from both the benign (blue) and pathogenic (red) test datasets. In purple contour lines, a kernel density of the simulated variant distribution is plotted.

